# Long-read nanopore shotgun eDNA sequencing for river biodiversity, pollution and environmental health monitoring

**DOI:** 10.1101/2024.11.01.621618

**Authors:** Orestis Nousias, Fiona G. Duffy, Isabelle J. Duffy, Jenny Whilde, David J. Duffy

**Affiliations:** The Whitney Laboratory for Marine Bioscience and Sea Turtle Hospital, University of Florida, St. Augustine, Florida 32080, USA; Department of Biostatistics, Yale School of Public Health, New Haven, CT, USA; Current address: Biotech Research and Innovation Center (BRIC), University of Copenhagen, Denmark; Department of Biology, College of Liberal Arts and Sciences, University of Florida, Gainesville, Florida, 32611, USA

## Abstract

As global temperatures rise, species populations and biodiversity decline, and infectious diseases emerge all at unprecedented rates, it is more vital than ever to accurately understand the current state of natural habitats. While traditional methods including direct sampling and observation have their merits, newer technologies offer additional sampling capabilities.

Here, we assess the feasibility of a single assay: shotgun long-read sequencing, to monitor species eDNA from across the tree of life, from viruses to complex multicellular organisms, across a river system (mountain tributary to sea). We conducted eDNA sampling and shotgun long-read sampling from water taken from the Avoca River watercourse, Co. Wicklow, Ireland, from a mountain tributary through to the sea, and comparative nearby beach sand sampling. We report that shotgun long-read sequencing and metagenomic analysis have utility for the detection and quantification of organismal DNA present in eDNA samples, from across the tree of life, from microbes (including DNA viruses) to mammals. With this single assay we were able to simultaneously quantify differences in eDNA abundance for a broad range of biodiversity and pathogens across sites and sample types. This included human, wildlife, plant and microbial pathogens and parasites with health, agricultural and economic importance. Additionally, the generated eDNA genomic data enabled population genetic applications even from natural complex community settings, as demonstrated here for blue mussels (*Mytilus edulis*). The results demonstrate that Oxford Nanopore sequencing provides a quantitative approach for river biodiversity, pollution and environmental health monitoring.

This method is more cost-effective and requires less laboratory preparation time or molecular expertise (barcode/primer design) than alternative biodiversity eDNA approaches (e.g. metabarcoding or qPCR). Shotgun analysis approaches will also benefit from the continual expansion of reference genome databases, for environmental, evolutionary and medical reasons, among others. Computation, cloud and artificial intelligence (AI) tools (such as the cloud-based analyses utilized here) can analyze shotgun sequencing eDNA data in a matter of hours and can be used by conservation practitioners, environmentalists and public health personnel without the need for coding or in-depth bioinformatics skills. The proven citizen/community scientist applicability and ease of eDNA sampling may further revolutionize and democratize biodiversity research, conservation surveillance, and environmental health monitoring. Long-read shotgun sequencing of eDNA offers the means to assess whole ecosystems, and the ecological, trophic, and host-pathogen interactions occurring within them.

## Introduction

Rapid advances in genomic sequencing technologies and computational analyses are presenting novel opportunities for non-invasive monitoring of environments, microbes and wild species, including human-impacted habitats. Such technologies are set to play an increasingly important role in studying and responding to the ongoing interrelated international crises of biodiversity loss, habitat loss, climate emergency, infectious disease epidemics and anthropogenic pollution of biomes^1^.

These technologies will be crucial for effective management, conservation and nature restoration, including meeting Target 3 of the Convention on Biological Diversity’s Kunming-Montreal Global Biodiversity Framework (GBF). As part of the global effort to halt and reverse biodiversity loss, this target includes the ambitious goal of increasing the global coverage of protected areas and implementing other effective area-based conservation measures to at least 30% by 2030 (referred to as ‘30×30’)^2–4^. To ensure the areas most capable of sustaining the richest biodiversity are protected, and to measure the success or otherwise of implemented protections, scalable, rapid, cost-effective tools are required for biodiversity assessment across differing regions of the globe with highly varied flora, fauna, funga, viromes and microbiomes. Advanced genomic sequencing platforms have been coupled with novel sampling approaches to recover genetic material from environments, providing new tools for ecosystem, pathogen and biodiversity monitoring and wildlife conservation ^5–18^.

Despite its ubiquity in fields such as human medical research, there is a relative paucity of studies applying whole genome (nuclear and organelle DNA) sequencing approaches to environmental samples, particularly for multicellular organisms and long-read sequencing technologies^13–15,19–22^. Whole genome sequencing approaches in which all genetic material present in a sample is sequenced without enrichment or targeting of specific regions is known as shotgun sequencing when applied to environmental samples. This paucity of genomic shotgun sequencing eDNA studies in the scientific literature occurs despite early comparisons revealing that metabarcoding is less consistent than shotgun whole mitogenome sequencing ^22^.

eDNA is beginning to be adopted as a non-invasive tool for biodiversity and pollution monitoring. For example, an innovative recent study demonstrated that eDNA content is correlated with long-term water quality indicators and riverine biodiversity measures^23,24^. However, to achieve broad taxonomic coverage, 14 separate metabarcoding assays were needed^23^, requiring significant laboratory manipulation and processing time. In contrast, single-assay shotgun sequencing is rapid, reduces barcode/primer and PCR bias, and therefore provides more directly comparative quantitative data across all recovered species groups. Additionally, it can recover species beyond those recoverable even with 14 metabarcoding assays, and importantly it recovers genomic data beyond the small barcode regions used by metabarcoding approaches. While some pioneering studies have successfully applied long-read sequencing to aquatic systems, these are normally confined to bacterial or fungal analyses and/or metabarcoding approaches^9,25–27^.

As reference genome databases are increasingly being populated with high-quality genomes of species from across the tree of life^3,28^, the species-level resolution of eDNA shotgun sequencing will continue to improve. Additionally, improved bioinformatic pipelines to filter out promiscuous reads will help to further increase the specificity of genus and species calls. Continued improvements in sequencing technologies, including higher output, will likely further enhance the already impressive sensitivity of such approaches. Cloud-based automated analysis workflows and artificial intelligence (AI) will improve the useability of such tools, making them more accessible to a wide range of stakeholders.

Here, we conducted eDNA sampling and shotgun long-read sampling from water samples taken from the Avoca River watercourse, Co. Wicklow, Ireland, from a mountain tributary of the river through to seawater samples near the river mouth (Supplemental Fig. 1). For comparison, we also collected and sequenced beach sand eDNA from 5 km north of the Avoca River mouth (Supplemental Fig. 1). We demonstrate the simultaneous detection of biodiversity from viruses to vertebrates in a single assay: long-read shotgun sequencing of eDNA samples. We reveal that this rapid single assay approach provides a viable complementary or alternative approach to traditional eDNA metabarcoding^14^. This approach could successfully generate results related to biodiversity, pollution and environmental health and viral monitoring, for a representative river system, from mountain tributary to the Irish Sea. As sequencing technologies and associated computational analyses continue to advance, long-read shotgun sequencing is poised to become a rapid monitoring tool for changing environments, including aquatic point source tracing of biotic pollution, and for disentangling the relative contribution of different biotic pollution sources (human waste, agricultural, industry or wildlife). Simultaneously, this biological ‘everything’ assay can monitor changes to biodiversity and species distributions in response to perturbations, including anthropogenic pollution, habitat destruction and beyond.

## Results

### Recovery of total DNA from environmental samples

We assessed the ability of a single assay: shotgun long-read sequencing, to monitor species eDNA from across the tree of life, from viruses to complex multicellular organisms, across a river system (Supplemental Fig. 1). Irish eDNA samples were collected for two purposes: i) to quantify the level of human eDNA (species-specific qPCR) present in each 2022 sample^19^, and ii) to assess the viability of long-read shotgun sequencing to monitor all detectible DNA per sample (ONT) (reported here). The highest eDNA yields were obtained from the tidal portion of the Avoca River (boat ramp and harbor samples) that runs through Arklow Town, Co. Wicklow (Fig. 1a, Supplemental Fig. 1a,b). Lower quantities of eDNA were recovered from the mountain stream and seawater sites, despite larger volumes being filtered (Fig. 1a, Supplemental Table 1). This indicates a higher abundance of biological material at the tidal regions of the river. Despite having lower total DNA abundances (Fig. 1a), the two sea water sites had similar eukaryotic (animals, plants, fungi and many unicellular organisms) eDNA concentrations to the harbor samples (pan-eukaryotic 18s rRNA gene qPCR assay^19^, Supplemental Fig. 2a).

**Figure 1.**
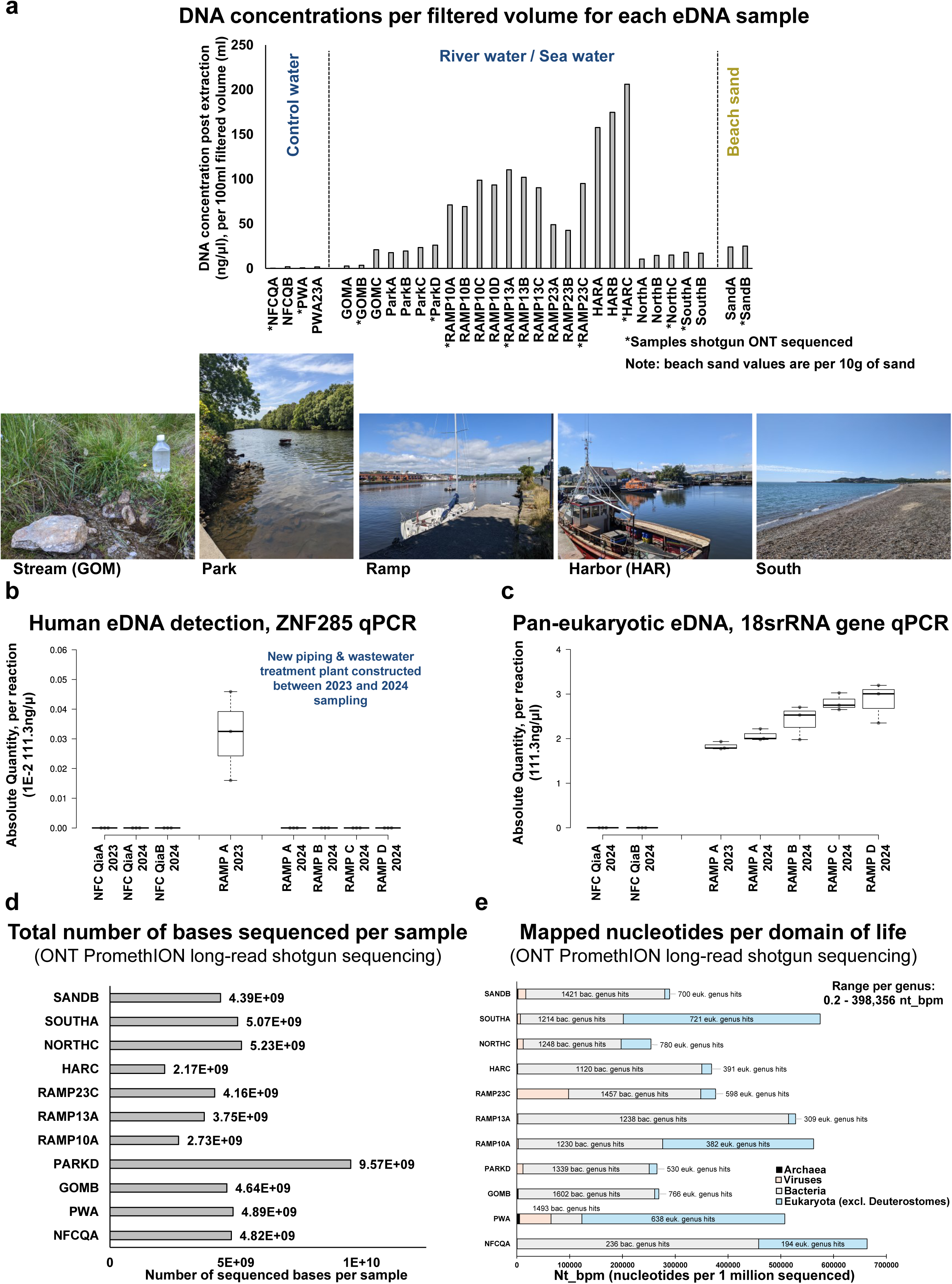
**a)** Graph of DNA concentrations as measured by Nanopdrop Spectrophotometer (Thermo Fisher) per 100 mls filtered volume for each eDNA sample, with images from the sampling sites inset. For total volumes filtered for each of the sequenced samples see Supplementary Table 1. For pan-eukaryotic eDNA concentrations per sample, see Supplementary Figure 2a. **b)** qPCR-based species-specific quantification of human eDNA from Avoca River water boat ramp sampling site (2023 and 2024). Quantified with ZNF285 human specific assay. Each qPCR reaction is a 10 μl reaction containing 1 μl of extracted eDNA template. Absolute quantity of human eDNA per sample was calculated by comparing with a ZNF285 assay on a human cell line (IMR32) DNA standard curve. **c)** Pan-eukaryotic eDNA levels within ramp site (2023 and 2024) water eDNA samples. Absolute quantity of 18 s rRNA pan-eukaryotic eDNA levels, as assessed by qPCR. Each qPCR reaction is a 10 μl reaction containing 1 μl of extracted eDNA template. Absolute quantity of pan-eukaryotic eDNA per sample was calculated by comparing with an 18s s rRNA pan-eukaryotic assay on a human cell line (IMR32) DNA standard curve. **d)** Graph of the total number of nucleotide bases sequenced per sample, by Oxford Nanopore Technologies (ONT) PromethION long-read shotgun sequencing. **e)** Graph of the number of mapped nucleotides per domain of life (by CZ ID analysis, post filtering and promiscuous read removal) recovered by shotgun long-read ONT sequencing, nucleotide bases per 1 million bases sequenced (nt_bpm). The number of bacterial and eukaryotic (excluding deuterostomes) genus hits are shown, in or adjacent to relevant domain boxes. Note: all genus hits are reported with no minimum nt_bpm cut-offs applied. The nt_bpm range per genus recovered was from 0.201244 nt_bpm (Philodina rotifers in the PARKD sample) to 398,356 nt_bpm (Bradyrhizobium bacteria DNA in the NFC sample).

### Human eDNA for temporal and spatial wastewater release/contamination monitoring

We previously showed by species-specific human qPCR of the 2022 samples (Supplemental 2b, Whitmore et al. 2023^15,19^), that the river is heavily contaminated by human eDNA at the boat ramp site in Arklow town. The majority of this human eDNA is suspected to originate from improperly treated Arklow town wastewater entering the river (https://www.epa.ie/publications/monitoring--assessment/waste-water/Urban-Waste-Water-Treatment-in-2022-Report.pdf). Sampling at the same site in 2023 confirmed the continued presence of human eDNA (Fig. 1b). However, in 2024 human eDNA was no longer present at detectable levels (Fig. 1b), despite high eukaryotic eDNA recovery rates (Fig. 1c). This suggests that the extensive wastewater treatment plant and pipe construction works undertaken in Arklow between 2023 and 2024 successfully removed/diverted this pollution source from this site.

### Pan-biodiversity assessments from a single assay: shotgun long-read sequencing

Of the 31 water and sand eDNA samples collected in 2022 (n = 25) and 2023 (n = 6), 11 representative samples were selected for long-read sequencing (Fig. 1a,d,e and Supplemental Table 1). These unbiased shotgun sequencing data provide insights into the relative abundance of eDNA recovered from each domain of life (Fig. 1e) and animal phylum (Fig. 2a). Bacterial reads were the most abundant in all samples except three: well water (PWA), river water (RAMP10A) and sea water (SOUTHA, Fig. 1e). For these three samples, eukaryotic DNA (excluding Deuterostomes) was most abundant (Fig. 1e). However, all samples including these three had more bacterial genera called than eukaryotic genera (excluding Deuterostomes), indicating that bacteria were the most diverse domain of life (Fig. 1e). Archaea were most abundant in private well ground water (5,224 nt_bpm), followed by beach sand (2,225 nt_bpm, Fig, 1e, Supplemental Fig. 3a). Across all environmental samples, the metazoan (animal) phylum with the highest read counts were Arthropods (Fig. 2a). Rotifer DNA was highest in the groundwater sample (private well pump, domestic water), while Chordata DNA was highest in the beach sand, followed closely by the northern sea water sample near the Avoca River’s outflow into the Irish Sea (Fig. 2a, Supplemental Fig. 1a). There was a mix of overlapping and unique metazoan species detected between the eDNA samples (Fig 2b, Supplemental Fig. 3b). No counts were recovered in any sample for the deep sea-restricted Xenoturbellida phylum^29,30^. With a minimum cut-off of five counts per species, 861 metazoan species were detected in 10 g of beach sand (SOUTHB, Fig. 2b). The sea water sample (NORTHC) had the second highest diversity of metazoan species detected, with 735 (Fig 2b, Supplemental Fig. 3b). The negative field control (Qiagen molecular grade water) had DNA from 24 metazoan species detected (Fig. 2b), while domestic drinking water from a private groundwater pump (near the mountain stream sampling site, PWA) had DNA from 281 metazoan species detected (Fig. 2b, Supplemental Fig. 3b). Species richness was not a factor of the number of bases generated per sample (Fig. 1d, Supplemental Table 1).

**Figure 2.**
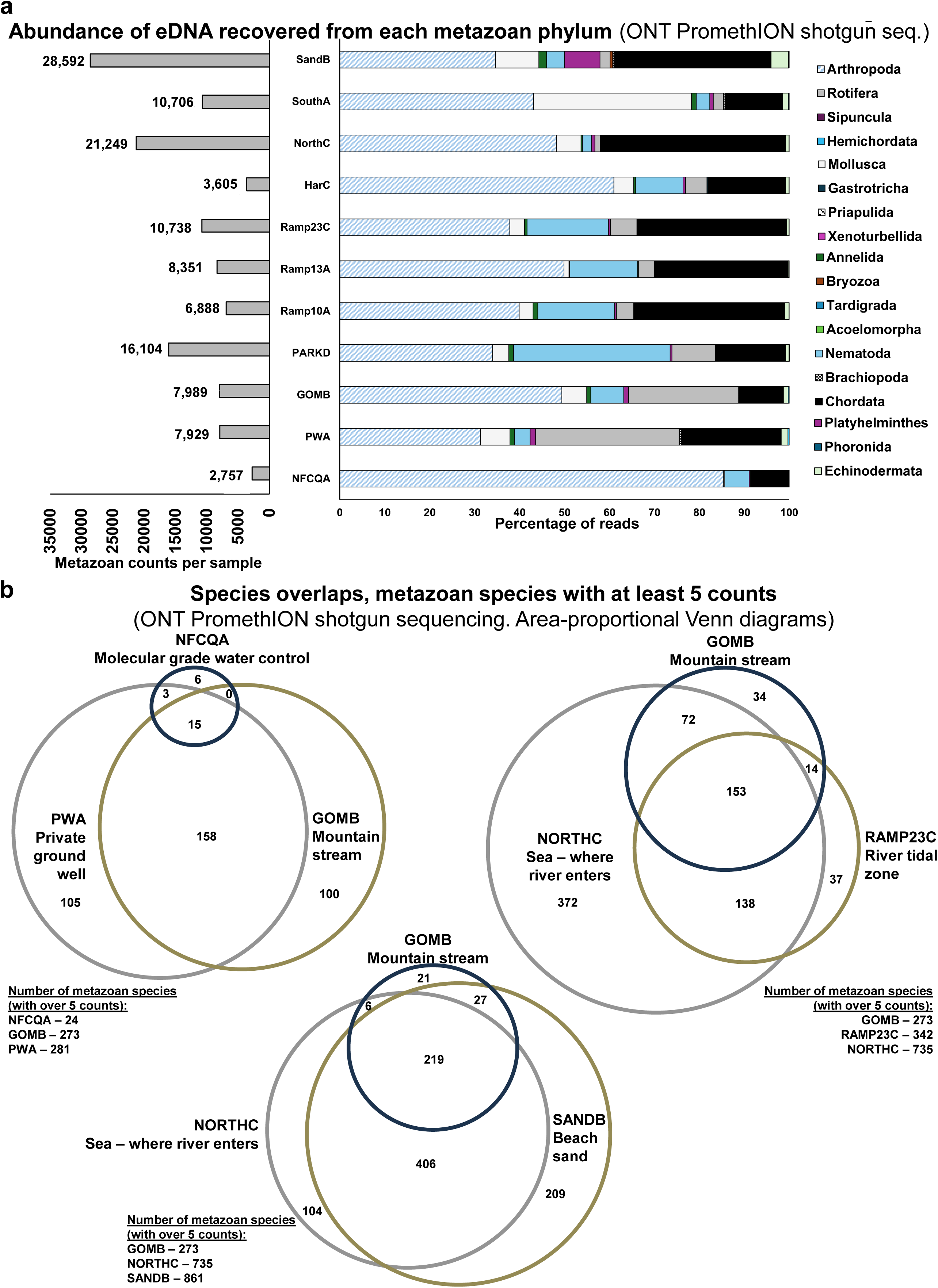
**a)** Abundance of eDNA recovered from each of the 18 metazoan phyla, by shotgun long-read ONT sequencing. *Left*: total metazoan counts per sample. *Right*: percentage of metazoan counts per phylum, per sample. Note: no counts were recovered, in any sample, for the deep sea Xenoturbellida phylum^29,30^. **b)** Area-proportional Venn diagrams of metazoan species overlaps between eDNA samples, showing the number and overlap of metazoan species with at least 5 counts per sample, as detected by ONT shotgun sequencing. Generated using BioVenn ^48^. See Supplemental Figure 3c for additional sample comparisons.

### Species eDNA abundance quantification and population genomics

We next compared the top 10 eukaryotic and non-eukaryotic hits in the beach sand (BEACHB) and tidal river water (RAMP23C) samples. For the top hits in both samples the amount of non-eukaryotic DNA in the environmental substrates was generally an order of magnitude greater than eukaryotic DNA (Fig. 3a). Eukaryotic organisms have cells with a membrane-bound nucleus (i.e., animals, plants, fungi, and many unicellular organisms). Our non-eukaryotic category includes bacteria, archaea and viruses. Note that this analysis omits deuterostomes (a sub-class of eukaryotes), which the CZ ID tool used filters out as potential host species^31^. The most abundant non-eukaryote in the river water sample was Asteriusvirus, from a family of bacterial and archaeal viruses (Fig. 3a). The most abundant non-eukaryote in the beach sand was the bacteria genus Mycobacterium, consisting mostly of *Mycobacterium branderi* (Fig. 3a). Seven of the water sample’s (RMAP23C) top 10 non-eukaryotic hits were bacteria; conversely, seven of the beach sand’s (SANDB) top 10 non-eukaryotic hits were viruses, including uncultured marine viruses (Fig. 3a,b).

**Figure 3.**
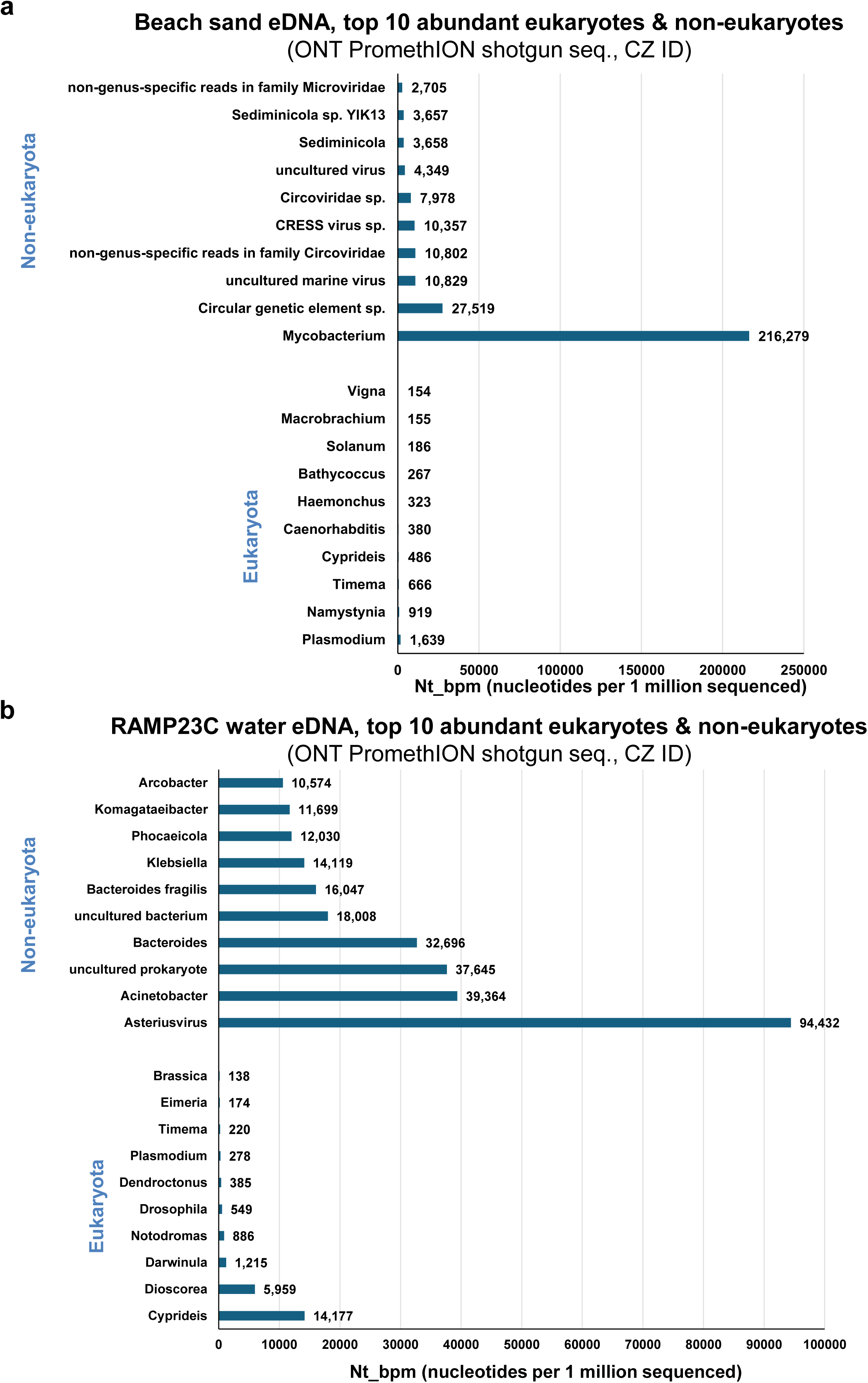
**a)** The top ten most abundant eukaryotic and non-eukaryotic genera eDNA (by nt_bpm) as detected by long-read sequencing and CZ ID analysis of the beach sand sample. **b)** The top ten most abundant eukaryotic and non-eukaryotic genera eDNA (by nt_bpm) as detected by long-read sequencing and CZ ID analysis of the Avoca River, Arklow Town (boat ramp) water sample RAMP23C.

Human DNA was also quantifiable by shotgun sequencing in environmental samples (Fig. 4a), with the highest eDNA levels being in Avoca River samples taken in Arklow town. This mirrors previous human eDNA assessment using species-specific qPCR assays; improperly treated wastewater release in Arklow is the suspected primary source of this human eDNA^19^. The boat ramp site sample (RMAP13A) from July 2022 had the highest level of human eDNA (2,279 counts), followed by the car park (PARKD, July 2022) and same boat site (RAMP23C, July 2023) sampled one year later (1,392 and 1,270 counts respectively). By applying shotgun sequencing to these samples, the abundance of human eDNA can be directly compared to that of other animals, human pathogens, and gut/fecal-associated microbes. We next compared the level of human eDNA recovered to that of a selected group of human-associated and wild mammal species (Fig. 4b). For this group of mammals, human eDNA was the most abundant in almost all samples, with the main exception being beach sand, which had a higher abundance of otter, rabbit, ferret and dog eDNA (Fig. 4b). In fact, of the 17 non-human mammal species analyzed, the beach sand had the highest abundance for 13 of these mammals, indicating a role of sand in filtering and accumulating eDNA. Only dog, cat, stoat and house mouse eDNA was more abundant in at least one of the water samples (Fig. 4b). The three highest abundances of eDNA recovered across these 17 mammals were dog, ferret and otter (137 to 98 counts). Across all 17 of these mammals the total count was highest in the sand sample (SANDB, 772 count), followed by a sea water sample (NORTHC, 534 count), while the negative field control and mountain stream only had small numbers of total counts for these mammals, 16 and 20 respectively.

**Figure 4.**
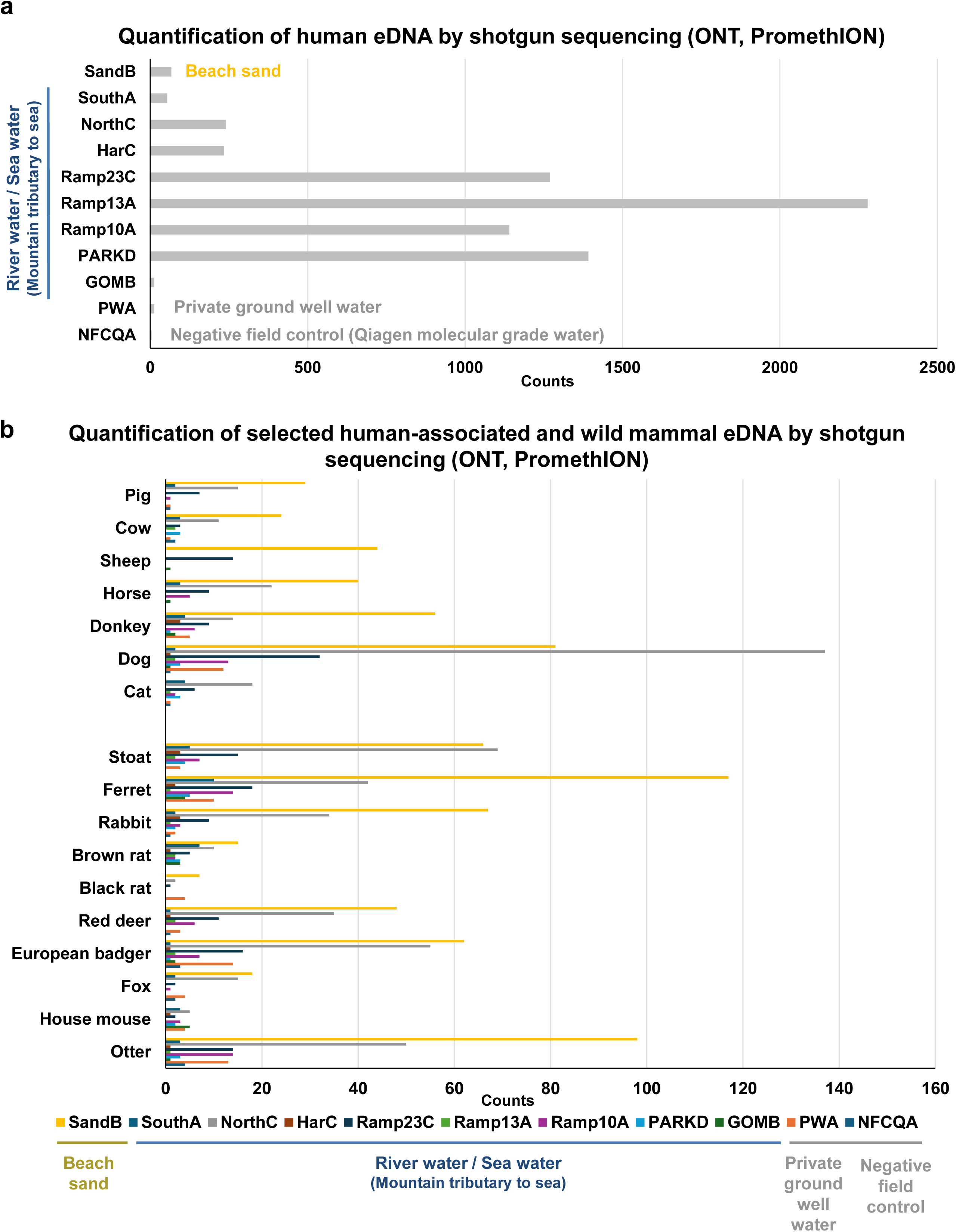
**a)** Quantification of human eDNA (read counts) across the sample set, as detected by metagenomic analysis of long-read shotgun sequencing. **b)** Quantification of selected human-associated mammals and wild mammals eDNA across the sample set, as detected by metagenomic analysis of long-read shotgun sequencing.

Next, to test the ability of shotgun sequencing long-read eDNA to determine the relative abundance of farmed species eDNA in waterways, we assessed detection of Oncorhynchus (fish group) eDNA. There is a rainbow trout (*Oncorhynchus mykiss*) fish farm approximately 8 km upstream of the boat ramp sampling site, and the mountain stream site (GOMB) is further upstream from the trout farm. No rainbow trout eDNA was detected in the mountain stream site (GOMB) (Fig. 5a). However, almost all downstream samples contained rainbow trout eDNA, with the highest abundance being the site closest to the farm (PARKD, 42 counts, Fig. 5a), approximately 8 km downstream from the farm. We then assessed the abundance of eDNA from select aquatic metazoan species across the samples. Once again, the beach sand contained the highest abundance of many of these species (10 out of 15 of the detected species), including common octopus, coral and leatherback sea turtle (Fig. 5b). Of these 15 species, blue mussel (*Mytilus edulis*) in the seawater sample (SOUTHA) had by far the highest abundance (837 counts, Fig. 5b). The *M. edulis* mitochondrial aligning reads (CZ, ID) from the SOUTHA sample shotgun ONT sequencing were blasted (NCBI, BLASTn) against *M. edulis* nucleotides, and a phylogenetic tree constructed (BLAST pairwise alignment, minimum evolution tree) (Fig. 5b, inset). The blue mussel eDNA recovered from the Irish Sea, Arklow (SOUTHA), clustered most closely to blue mussel tissue DNA samples from Germany (Langeoog, Wadden Sea) and Wales (Swansea Bay, Irish Sea). This demonstrates that shotgun long-read sequencing can be utilized not just for species detection, but with sufficient eDNA recovery it can be used for non-invasive population genetics .

**Figure 5.**
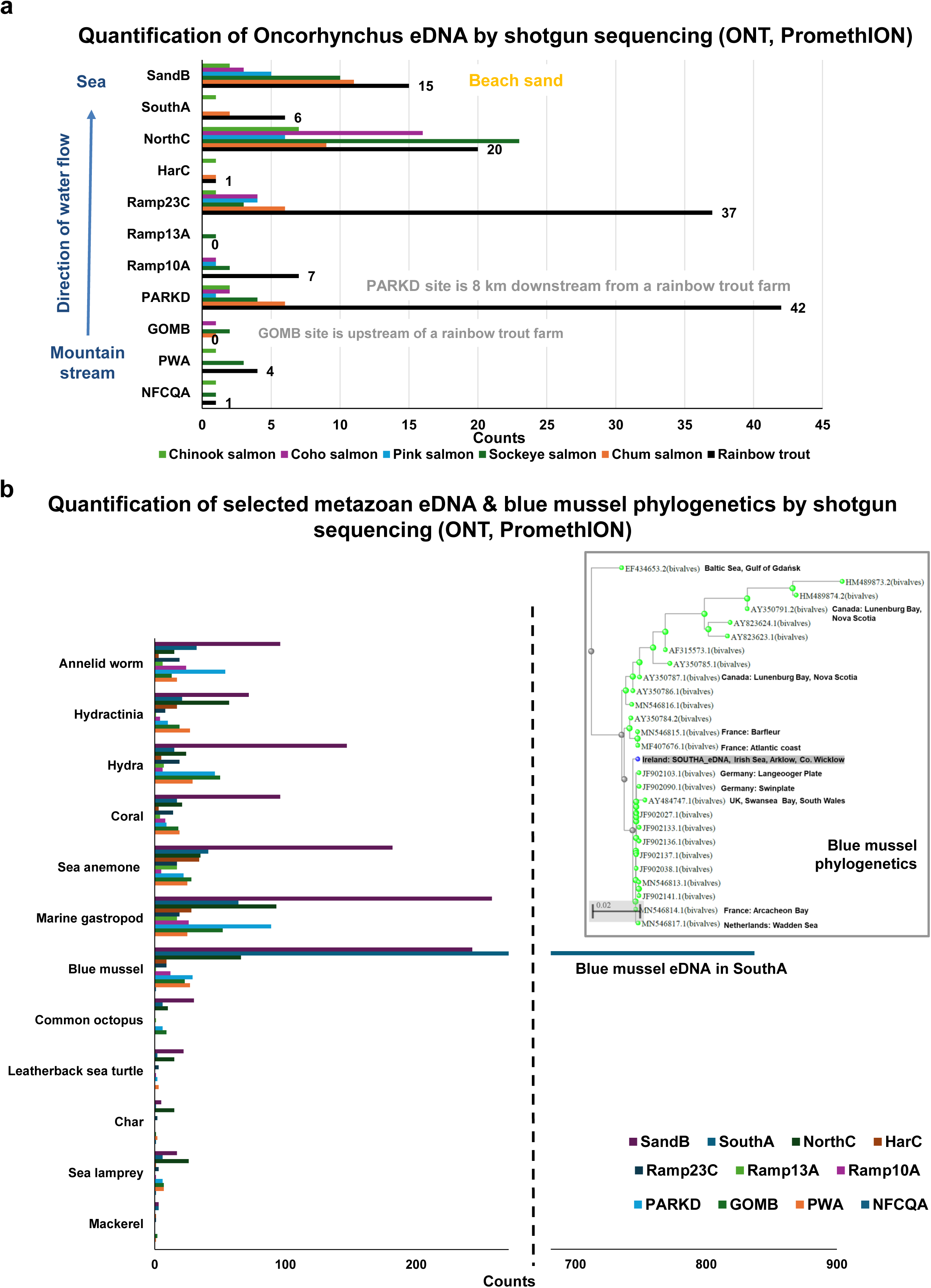
**a)** Quantification of Oncorhynchus (salmon/trout) species eDNA across the sample set, as detected by metagenomic analysis of long-read shotgun sequencing. **b)** Quantification of selected aquatic or aquatic sediment (Capitella, worm) metazoan eDNA across the sample set, as detected by metagenomic analysis of long-read shotgun sequencing. Insert: phylogenetic placement of the detected blue mussel (*Mytilus edulis*) eDNA, generated by NCBI Blastn analysis of the CZ ID identified *M. edulis* eDNA sequence. NCBI accession numbers of the Blastn retrieved comparative sequences are provided at the start of each branch name.

As expected, marine algae and plankton species also had highest abundances in the seawater samples (Fig 6a,b). Tree, flowering plant, butterfly and Trichoplax eDNA were also detected, at varying quantities between the samples (Fig. 6c). Human, animal and plant pathogens and parasites were also detected across the sample set (Fig. 6b,c). This includes the causative agent of global amphibian declines, the pathogenic chytrid fungus *Batrachochytrium dendrobatidis (Bd)*^32,33^ (Fig. 6c,d). The highest abundance of Bd was from the mountain stream sample, which drains upland bog and conifer plantations, prime amphibian habitat (Fig. 6d). This stream sample also contained the highest level of pine tree eDNA detected (Fig. 6c), and amphibian Ranavirus (2.82 nt_bpm). Of the river samples, this site also had the highest abundance of common frog eDNA, which is one of Ireland’s only three amphibian species (Fig. 6d). This result highlights the utility of rapid broadscale environmental screening for organisms, pathogens and parasites, which could then be followed up with more targeted monitoring of species of concern (species-specific qPCR/dPCR assays, etc.). Targeted study design and site selection could be greatly informed by the rapid generation of exploratory genomics eDNA data. This also highlights the power of eDNA genomics for data-driven surveillance and discovery, being able to quantify both pathogen and host without *a priori* knowledge of which pathogens are present at the sampling site.

**Figure 6.**
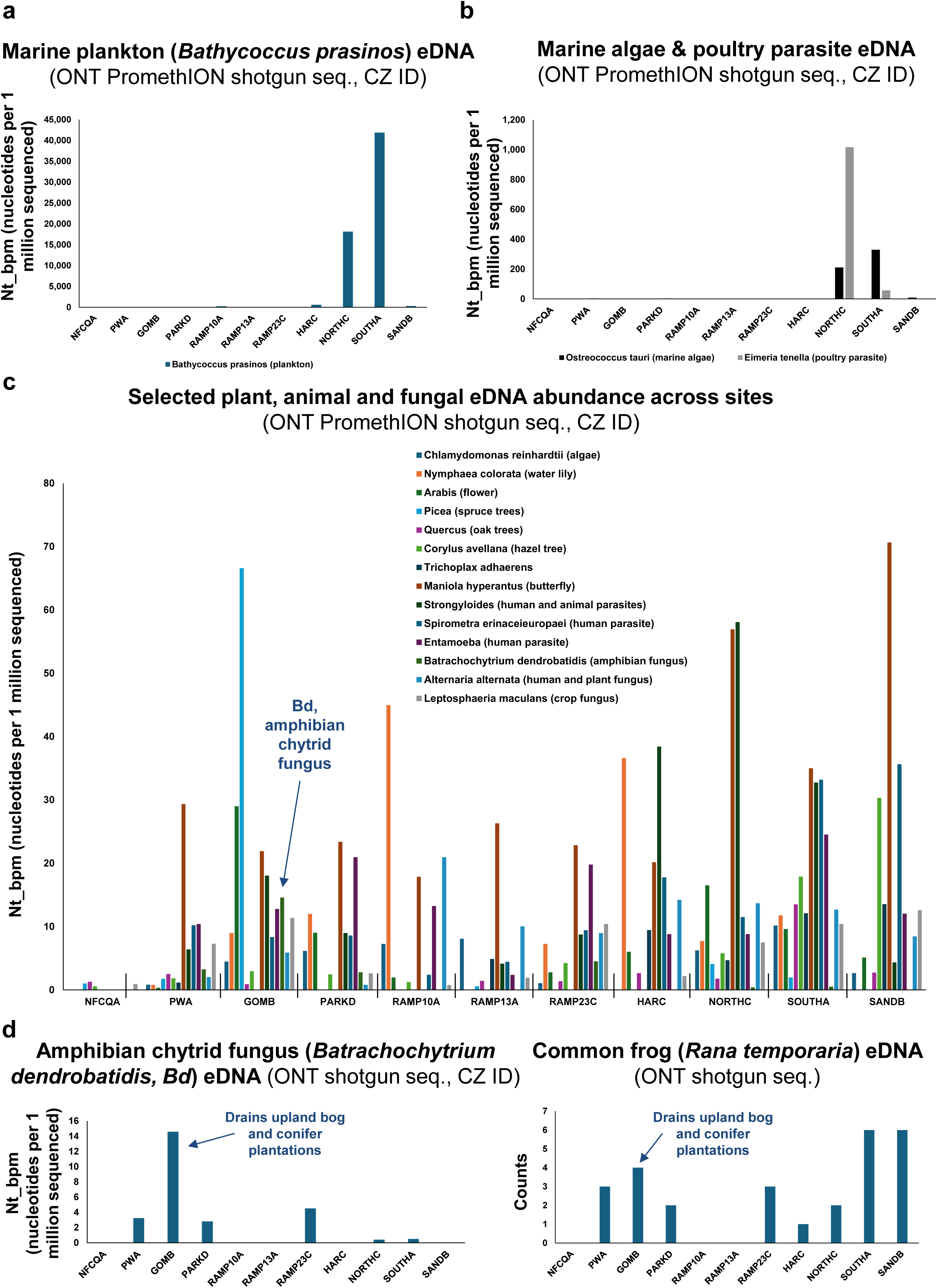
**a)** Quantification of the marine plankton *Bathycoccus prasinos,* throughout the sample set, arranged from freshwater to sea water (and finally beach sand), as detected by CZ ID metagenomic analysis of the long-read shotgun sequencing. **b)** Quantification of a marine algae (*Ostreococcus tauri*) and poultry parasite (*Eimeria tenella*) eDNA, throughout the sample set, arranged from freshwater to sea water (and finally beach sand), as detected by CZ ID metagenomic analysis of the long-read shotgun sequencing. **c)** Quantification of selected plant, animal and fungal species eDNA across the sample set, as detected by metagenomic analysis of long-read shotgun sequencing. The list includes parasitic species. **d)** Left: quantification of the amphibian chytrid disease agent *Batrachochytrium dendrobatidis* throughout the sample set, as detected by CZ ID metagenomic analysis. Right: quantification of the common frog (*Rana temporaria*) throughout the sample set, as detected by metagenomic analysis.

### Simultaneous quantification of pathogen DNA from eDNA genomics

Shotgun sequencing also recovered DNA from a variety of pathogens (Fig. 1e, 3a,b). We next assessed the total number of viruses detected in the beach sand (SANDB) and tidal river water sample (RAMP23C). Both samples had over 300 viruses present, with only 112 of these occurring in both samples (Fig. 7a). DNA from only 15 viruses was detected in the negative field control sample. The control sample also had much lower abundance of viral DNA recovered (range 225 - 0.64 nt-bpm) than the sand (range 27,519 - 0.23 nt-bpm) or river water sample (range 94,432 - 0.24 nt-bpm). We next examined each of these beach sand and river water samples for selected common human sexually transmitted skin, urinary or genital tract infection pathogens (Fig. 7b). We focused on these pathogens as they are some of the most likely to enter human wastewater systems. Diseases associated with these known human pathogens include genital herpes, periodontitis, urethritis, cervicitis, vaginitis and vulvovaginal candidiasis, gonorrhoeae, bacterial vaginosis and Legionnaires’ disease. Note, unlike conventional wastewater screening for human pathogens, our results come from environmental samples (beach and river) after DNA has left any wastewater treatment facility and mixed into environmental substrates containing complex eDNA mixtures. Despite this dilution effect “uncultured human fecal virus” was detected in both river water (RAMP23C, 503.26 nt-bpm) and beach sand (SANDB, 1.59 nt-bpm) samples. Seven of the beach sand’s (SANDB) top 10 non-eukaryotic hits were viruses, including uncultured marine viruses (Fig. 3a,b). Human- and animal-infecting pathogen and parasite eDNA from different domains of life, such as poxvirus, streptococcus and cryptosporidium could also be readily quantified from the data (Fig. 8a-c). Archaea were also readily detected by this approach, including human-gut- and acidic soil-associated species (Fig. 8d).

**Figure 7.**
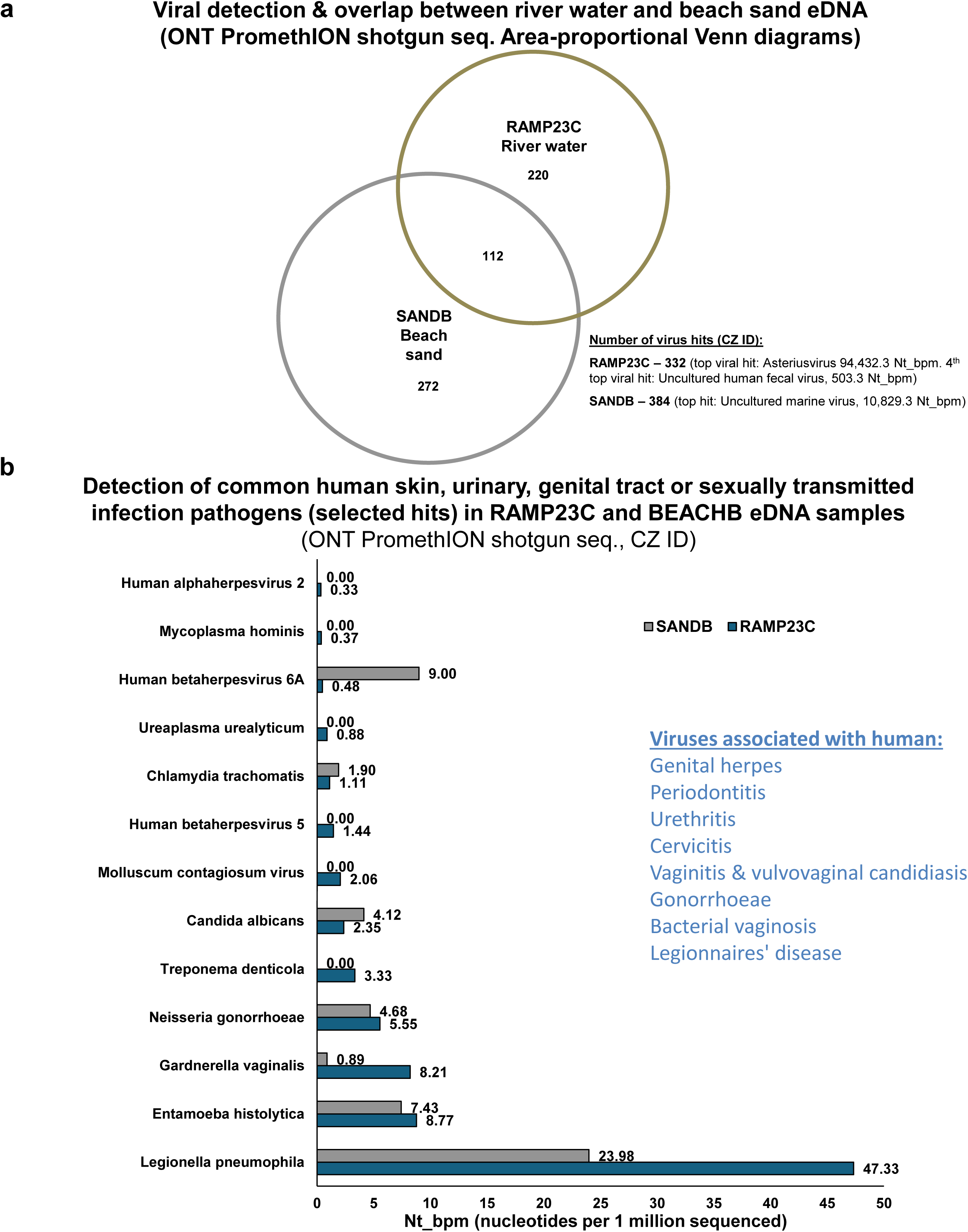
**a)** Area-proportional Venn diagrams of viral species overlaps between the beach sand and Arklow Town Avoca River water (RAMP23A) eDNA samples, as detected by CZ ID metagenomic analysis of long-read shotgun sequencing. Venn diagram was generated using BioVenn^48^. **b)** Quantification of common human sexually transmitted, skin, urinary or genital tract infection pathogen eDNA in the beach sand and Arklow Town Avoca River water (RAMP23A) eDNA samples. Note that at the time of sampling this river water sampling site was known to be contaminated with untreated human wastewater (Figs 1b, 4a and Supplemental Fig. 2b).

**Figure 8.**
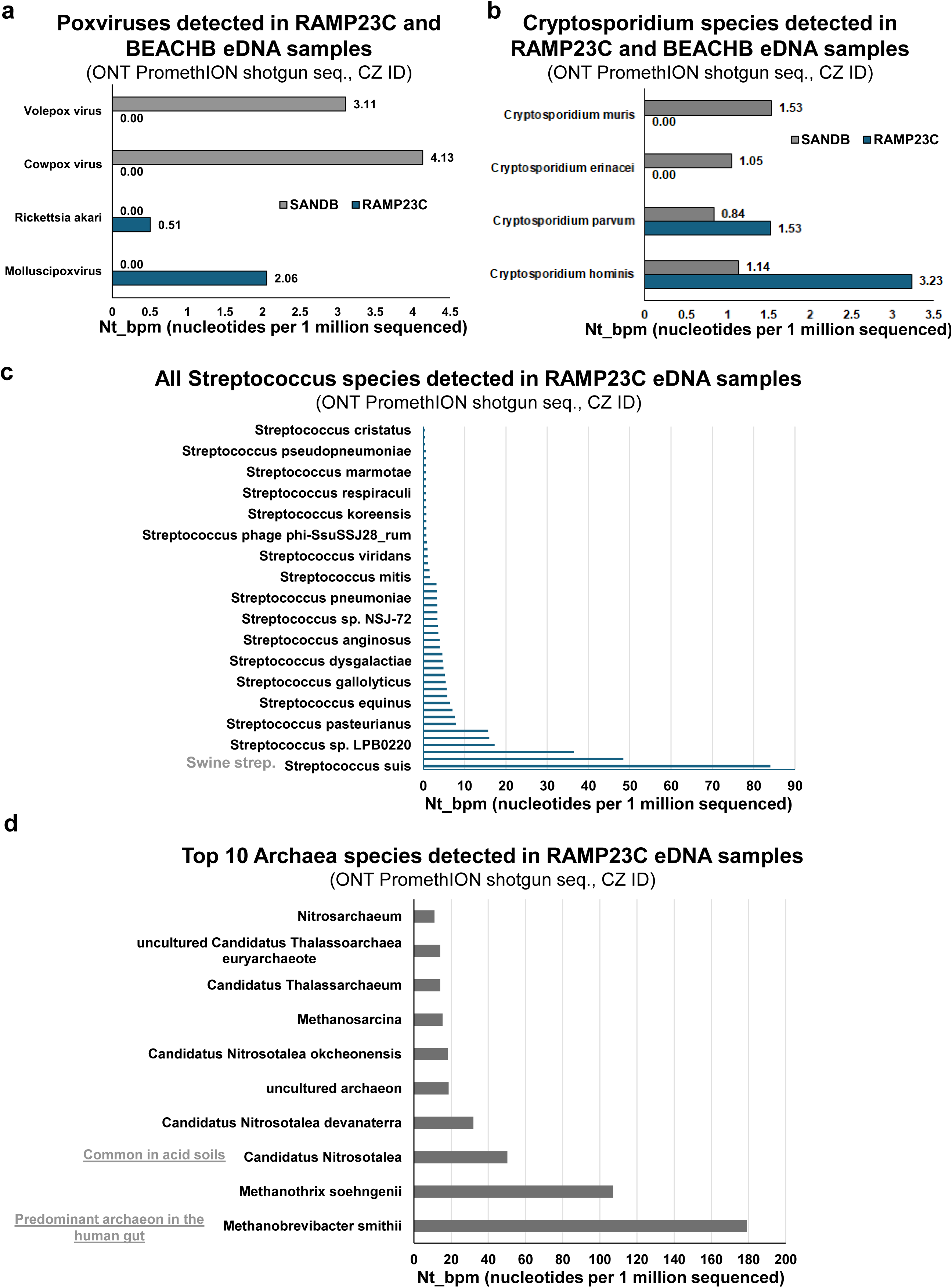
**a)** Quantification of poxviruses eDNA detected in the beach sand and Arklow Town Avoca River water (RAMP23A) eDNA samples by CZ ID metagenomic analysis of long-read shotgun sequencing. **b)** Quantification of Cryptosporidium (respiratory and gastrointestinal parasites) eDNA detected in the beach sand and Arklow Town Avoca River water (RAMP23A) eDNA samples. Detected by CZ ID metagenomic analysis of long-read shotgun sequencing. **c)** Quantification of Streptococcus bacteria species eDNA detected in the Arklow Town Avoca River water (RAMP23A) eDNA sample, by CZ ID metagenomic analysis of long-read shotgun sequencing. **d)** Quantification of the top ten Archaea (domain of single-celled organisms) species in the Arklow Town Avoca River water (RAMP23A) sample, as ranked by eDNA abundance. Detected by CZ ID metagenomic analysis of long-read shotgun sequencing.

Taken together, shotgun sequencing and identification of human eDNA and fecal and intestinal-associated microbe DNA offers a powerful tool for the point source identification of human wastewater entering waterways and aquatic bodies. Simultaneously, the information contained within shotgun sequenced eDNA samples enables ecosystem health analyses, biodiversity assessments and even monitoring of human, agricultural and biodiversity pathogen occurrence and abundance. Therefore, eDNA genomics offers cost-effective, non-invasive, population-level insights from environmental samples, to help address a range of societal and environmental issues.

## Discussion

Despite the early and innovative application of environmental genome shotgun sequencing to microbial eDNA of the Sargasso Sea in 2004 by Venter et al.^34^, the adoption of shotgun sequencing approaches to eDNA research has been surprisingly slow, particularly for the investigation of multi-cellular species. This apparent hesitation persists despite the interceding genomics boom of the last 20 years resulting in dramatic reductions in time and cost associated with shotgun sequencing, and simultaneously increases in ease, output and speed of sequencing and bioinformatic analysis. There has been a precipitous decline in the cost of deep sequencing, from billions of dollars for one human genome in 2000, to less than $200 today, making shotgun sequencing of eDNA ever more feasible, even for projects with limited budgets. The pace of advance of genomic technologies has not slowed, being largely driven by medical genomic applications^35^. The development of long-read sequencing should prove particularly beneficial to eDNA applications^20^. Environmental DNA for multicellular species has been dominated by less technologically-advanced metabarcoding approaches. It is surely now time to re-introduce and re-evaluate the role of modern shotgun sequencing technologies in transforming eDNA research capabilities.

We demonstrated here that long-read shotgun sequencing can be used as a single assay for simultaneous investigation of biodiversity, pollution, environmental health and viral monitoring, for an example river system from mountain tributary to the sea. This approach will automatically benefit from ongoing drives to increase high-quality reference genomes for eukaryotic and microbial species, as these new references are added to public databases. For example, 1.5 million eukaryotic reference sequence genomes are being generated by the Earth BioGenome Project (EBP) alone^3,28^. These approaches will also benefit from continued advances in the accuracy, speed and portability of long-read sequencing technology^3,20,25^. The data reported here showed good agreement between the shotgun data species-specific qPCR and known species trends, e.g. freshwater to marine transitions. However, not all species/genus calls in this proof-of-principle project may be fully correct, due to the current uneven coverage of species reference genomes in databases. For example, if a species does not yet have a reference genome, some reads may align instead to its closest-related species present in the queried database, rather than the actual species from which the DNA originates. However, this will improve over time in line with large-scale reference genome generation projects (see above). Similar issues arise with metabarcoding data, as barcode databases are far from complete, even for many charismatic specices^36–38^. Generally, for metabarcoding, the wider the species net is cast the less well-resolved the hits. Many metabarcoding studies and barcode sets do not achieve species-level resolution, opting rather to report at genus or higher level operational taxonomic units (OTUs).

The nature of shotgun sequencing data means that they can readily be reanalyzed against updated databases in the future, and with further refined bioinformatic pipelines, without requiring any additional laboratory work. Metabarcoding does not enjoy this benefit, as there are limitations to the depth and breadth achievable with a particular barcode set^36–38^. Sufficient sequence differences within the barcode may just not exist, even as more species are added to barcode databases, and any updated barcode sets would require re-PCR with the new barcodes and re-metabarcode sequencing of samples. Furthermore, shotgun sequencing is less susceptible to barcode and PCR biases^38^, and has much more tolerance of unknown genetic variation within populations, including distant populations from different locations around the globe^14^. Long-read shotgun sequencing also does not require specific advanced computational and laboratory effort to develop barcodes or qPCR assays. Additionally, it is becoming cost effective to sequence deeply and filter out promiscuous reads (reads aligning to multiple locations/genomes)^31,39^. The cost-effectiveness of sequencing and the high performance of computing makes this approach rival barcoding in terms of specificity, as only highly stringent reads can be analyzed. Generating additional information is a boon towards further research beyond just species identification.

We show here that in a single cost-effective and non-labor-intensive assay, rich information from across the tree of life can be recovered. In each environmental sample, with a single assay we recovered DNA from all domains of life from viruses to vertebrates. This approach is unbiased in revealing which groups contribute the most to eDNA abundance in any given sample. Direct comparison of eDNA quantities of different groups and domains of life is not readily achievable with barcoding- or qPCR-based approaches, due to the heterogeneity introduced by the different barcode/primer sets required for each group, and their differing efficiencies and affinities. The only domain of life not recovered by the shotgun DNA sequencing is RNA-based viruses. However, eRNA approaches coupled to shotgun sequencing (transcriptomics), either utilizing reverse transcribed cDNA, or direct long-read ONT RNA sequencing, can be used to recover RNA viruses from environmental samples. For example, SARS-CoV-2 has already been detected from wastewater^10,40,41^.

Fecal bacteria have long been used as a proxy in the study of sewage discharge, and, more recently, eDNA approaches have been adopted to target fecal bacteria^42,43^. Here, we show that in a single assay (shotgun long-read sequencing), human eDNA, fecal-associated bacteria, fecal-associated archaea and fecal-associated viruses can be simultaneously detected at a site with known frequent wastewater discharge. In 2022, Arklow was one of the 15 areas in Ireland that did not meet European Union wastewater treatment standards and was also one of the 26 Irish towns and villages discharging raw sewage into the environment every day (https://www.epa.ie/publications/monitoring--assessment/waste-water/Urban-Waste-Water-Treatment-in-2022-Report.pdf). In 2024, pipe improvement and wastewater treatment plant construction had progressed in Arklow town. The fact that human eDNA was no longer detectable (by species-specific qPCR) at the site (https://www.water.ie/projects/local-projects/arklow-wwtp/) highlights that infrastructure investment can effectively remediate some of the environmental damage caused by human wastewater. These eDNA results highlight the utility of human eDNA analysis to detect, monitor and point-source locate human wastewater pollution in aquatic systems, and to quantify improvements of remediation works.

Due to their ease of use, unbiased nature and scalability, long-read shotgun sequencing approaches can be used for widespread largescale biodiversity, pathogen or pollution screening/surveillance, with key results then being followed up with targeted species-specific approaches, such as qPCR/dPCR. Additionally, as shotgun approaches recover genetic information from across the genome (and mitogenome), the data generated can be applied to additional research areas such as population genetics^14,19^, as demonstrated here for blue mussels (*M. edulis*). Barcodes only recover information from informative, but small, restricted genetic regions, typically smaller than 200 bp. Shotgun eDNA sequencing enables a plethora of downstream analyses beyond just species identification, including viral variant analysis, population genetics, human genetic ancestry, and disease-associated variant calling^11,14,18,19^. Additionally, by simultaneously recovering both nuclear and mitochondrial genetic information, shotgun sequencing avoids biases in the temporal stability of mitochondrial versus nuclear eDNA^15^. This is an important caveat, not considered by the majority of animal/eukaryotic eDNA studies which predominantly rely on mitochondrial barcodes and mitochondrial qPCR/dPCR targets^14,15,44^. Indeed, these differences in nuclear to mitochondrial DNA degradation rates will likely prove to have utility for temporally discriminating the duration since eDNA release from its host species^15^.

Of all shotgun-sequenced eDNA samples in this study, beach sand had amongst the highest number of genera detected, and had a diverse metazoan phylum profile. This mirrors a similar dataset from Florida^15^, which indicated that beach sand acts as a natural filter of sea water, accumulating DNA from a wide variety of marine and terrestrial species (likely transported to the ocean by freshwater and air inputs). Therefore, we recommend detailed follow-up studies investigating the utility of beach sand sampling for robust marine (and terrestrial) biodiversity and species distribution surveying.

Environmental DNA research is reaching a crossroads. While metabarcoding approaches have proven incredibly powerful and have served the field well in establishing it as a strong force for biodiversity, pathogen and wildlife monitoring, metabarcoding only scratches the surface of the rich genomic data present in environmental samples. eDNA can be used to uncover secrets hidden in those parts of genomes that sit outside the limited short barcode region. The pending transition towards more powerful sequencing approaches is akin to the transition that occurred in the medical field from microarrays (DNA/RNA) to next-generation sequencing (genomics/transcriptomics). Despite their unquestionable utility and significance in advancing biomedical research, microarrays were ultimately supplanted by more powerful and less biased shotgun (whole genome) sequencing. This change was initially delayed by investigators’ reliance on and trust of microarrays, due to years of use, investment, and more established experience in the associated lab and analytical techniques. At its peak, 11,429 microarray papers were published in 2012, which dropped to 3,480 papers in 2024 (PubMed search for “microarray”, 01-Nov-2024). Conversely, in 2012 there were already 72,938 genomics papers, with 86,678 in 2024 (PubMed search for “genomics”, 01-Nov-2024). Eventually, more experience and trust in new sequencing technologies was built as these technologies were demonstrated again and again to outperform older microarray approaches, and the entire field transitioned to sequencing technologies (including RNA-seq). Having experienced that technological transition firsthand, we suspect that we are on the cusp of a similar technological revolution with shotgun sequencing in the field of eDNA. We also suspect that a similar pattern will play out: initial resistance and skepticism followed by growing acceptance, and then a near-ubiquitous switch.

This change will be aided by the versatility, continued reduction in cost, reduced labor and laboratory time, increased portability, improved analytical pipelines, and richer data provided by shotgun sequencing^3,13–15,19–22,34^. This paper aims to contribute to that discussion and adoption, by highlighting the extensive utility of applying shotgun long-read sequencing approaches to complex natural community eDNA samples.

### Summary

Here we demonstrate the feasibility of applying long-read shotgun sequencing to aquatic and sand eDNA samples, and its ability to recover genomic information from across the tree of life. We show how this information can be utilized to generate an unbiased quantification of the abundance of DNA between different domains of life, phyla, genera and species. We also demonstrate that, for species with sufficient DNA abundance in the sample, analyses beyond mere species identification are possible. In this case, population genetics from complex natural mixed community aquatic eDNA samples were carried out, without requiring targeting or amplification of specific DNA targets. Therefore, we call for increased adoption and analysis of shotgun sequencing for eDNA applications (including further verification and testing of its benefits and limitations). Such adoption should not be restricted to microbial life, but used for the simultaneous study of all domains of life. We also call for targeted investment (funding and research effort) to develop, optimize and make available eDNA shotgun sequencing analysis tools, with an emphasis on ease of access and usability. We stand at a crossroads where, rather than resistance to shotgun sequencing approaches for multicellular eDNA applications, adoption, application and rigorous assessments should be embraced. Shotgun sequencing will, at a minimum, complement the current dominant eDNA approaches, and in all likelihood exceed the capabilities of amplicon-based approaches, succeeding them as the most frequently applied eDNA tool in the near future. This approach is poised to enable biodiversity and wildlife assessments on the scale required to successfully implement and monitor the Convention on Biological Diversity’s GBF ‘30×30’ area-conservation targets.

## Methods

### Sample collection, DNA extraction and sequencing

All laboratory procedures from sampling through to final analysis (sequencing or qPCR) were conducted in a way that minimized any human DNA contamination, including any human DNA contamination from investigators. This included no contact of investigators/samplers with substrates being sampled (water or sand), new nitrile gloves being used for each sample collection, and frequent glove changes throughout sample processing, cleaning of equipment and benchtops with bleach prior to use, and negative field controls being treated identically to genuine field samples throughout all processes from extraction through to qPCR/sequencing (see below). All eDNA samples were extracted in a chick/mouse developmental biology lab that does not process human samples: the Murphy Lab, Zoology Department, Trinity College Dublin, Ireland.

River, estuarine, sea water and beach sand samples were collected between 2022 and 2024, on summer fieldwork visits to Co. Wicklow, Ireland. Negative field control samples of 500 ml Qiagen Nuclease-free water (Cat. No. 129117) was transported from the laboratory to environmental sampling locations and stored in a cool box with the environmental samples to monitor for potential contamination during sampling, transportation and processing. Negative field controls were filtered and extracted alongside the other collected sand and water samples from each sampling trip and subjected to the same next generation sequencing conditions and qPCR conditions. A range of 180 ml - 600 ml water was filtered per sample (until the filter was clogged with material, or a maximum of 600 ml was reached) (see Supplemental Table 1 and Whitmore et al., 2023^19^). Filtration was performed shortly after sampling using 0.22 µm pore Millipore Sterivex-GP Pressure Filter Units (Merk Millipore Cat No: SVGPL10RC) for each DNA sample^11,14^. Water samples were pumped (by hand) through 0.22 µm Sterivex-GP Pressure Filter Units and capped with B.Braun luer lock caps (Medline Cat No: BMGTMR2000B). Hand pumping was carried out with sterile 50 mL BD luer lock syringes (Fisher Scientific, 13-689-8 BD). 740 μl of Qiagen Buffer ATL was then added per filter and the filters stored at -20°C until transport to the lab (Trinity College Dublin) and DNA extraction. For sand eDNA, a 50 ml tube was filled with sand from each sampling event, with 10 ml of this sand used per individual eDNA extraction^14^. Sand was stored at -20°C until transported to the lab (Trinity College Dublin).

For water samples, filters were thawed and 60 µl Proteinase K from a Qiagen DNeasy Blood and Tissue Kit were added to each sample (filter) and they were placed in sterile 50 mL Falcon conical centrifuge tubes in a rolling incubator overnight at 56°C.

For sand samples, Qiagen Nuclease-free water (Cat. No. 129117) was added to each individual 10 ml sand sample in individual 50 mL Falcon conical centrifuge tubes, at approximately two times the volume of sand (10 mL sand and 20 mL water). Samples were shaken gently by hand then set on a rocking platform for 1 hour at room temperature, with additional gentle shaking by hand every 15 minutes. Samples were then rested until sand had sunk to the bottom of each tube, then the supernatant was immediately pipetted into a 50 mL sterile BD luer lock syringe (Fisher Scientific Cat No: 136898). Samples were then hand filtered using 50 mL BD luer lock syringes through 0.22 µm Sterivex-GP Pressure Filter Units (Millipore Cat No: SVGPL10RC) and capped with B.Braun luer lock caps (Medline Cat No: BMGTMR2000B). 740 µl Buffer ATL and 60 µl Proteinase K from a Qiagen DNeasy Blood and Tissue Kit (Qiagen Cat No: 69504) were added to each sample and they were placed in sterile 50 mL Falcon conical centrifuge tubes in a rolling incubator overnight at 56 °C. For both water and sand samples, after overnight incubation the rest of our eDNA extraction method using a modified Qiagen DNeasy Blood and Tissue kit (Qiagen, Cat. No. 69504) protocol^11,14,15,19,45^ was applied. After overnight incubation, solutions were transferred from the Sterivex-GP Pressure Filter Units to 2 mL microcentrifuge tubes using 1 mL Fisherbrand™ Sterile Syringes (Fisher Scientific 14-955-462). Equal volume AL Buffer and ice-cold ethanol were added to each sample and they were vortexed and centrifuged after each addition, according to the rest of the kit manufacturer’s protocol, with the following exception: DNA was eluted with 70 µl AE Buffer (incubated at 70 °C prior to addition to spin column), incubated on the column at room temperature for 7 minutes and centrifuged into a 1.5 mL microcentrifuge tube at 6000xg for 1 minute. DNA concentration was measured on a ThermoScientific Nanodrop 2000 Spectrophotometer (at Systems Biology Ireland, University College Dublin) and samples were stored at -20 °C until qPCR or shotgun sequencing. Prior to filtration and between every sample, laboratory surfaces and equipment were cleaned with 70% ethanol and then 10% bleach (to destroy DNA). Collection bottles were disinfected (washed thoroughly) with 10% bleach and rinsed thoroughly with deionized water.

Previously extracted^46^ IMR32 gDNA, extracted using the standard cell line protocol of a Qiagen DNeasy Blood and Tissue kit (Qiagen, Cat. No. 69504), was used as template for generating standard curves (both human and pan-eukaryotic standard curves). This DNA had been extracted from cultured human IMR32 neuroblastoma cells in May 2013 at Systems Biology Ireland, UCD for the Duffy et al. (2018) study^46^, and had subsequently been stored at -20°C until used in this study (2024).

### Bioinformatics

Samples were shotgun sequenced commercially at BMK Gene, with Oxford Nanopore Technologies (ONT) libraries sequenced on a PromethION 48. All sequencing libraries were pooled equally (including NFCs) to give equal sequencing depth opportunity. All bioinformatic tools were utilized using default parameters, unless otherwise stated.

Chan Zuckerberg (CZ) ID^31,39^, a free, cloud-based metagenomics platform (https://czid.org/) was utilized for biodiversity bioinformatic analysis. The CZ ID platform queries genetic information from all species, excluding Deuterostomes. The CZ ID nanopore metagenomic pipeline was utilized. As the platform was primarily developed for microbial metagenomics it does not query deuterostome sequences as these are identified as host species, and it actively subtracts host reads such as human reads. Additionally, for samples with over 1M reads, the pipeline actively sub-samples only 1M reads per analysis. OTUs from the CZ ID data were identified twice: those identified with a single sequence, and those identified by at least two. To conduct metazoan metagenomics, including Deuterostomes, the CZ ID nanopore metagenomic pipeline was recreated in an HPC environment using bash and python, without host filtering, without sub-sampling to 1M reads, and aligning to all metazoan species (NCBI Blast). To classify metagenomic reads based on taxonomy, DIAMOND v2.1.7^47^ was utilized for its inherent high-speed alignment, sensitivity and the translation of nucleotide reads for protein alignments capabilities. Firstly, metagenomic reads from eDNA samples were pre-processed, for adapter trimming and quality control using Porechop v0.2.4^59^. The two-step DIAMOND process involved database preparation and read alignment. A reference database was constructed using the non-redundant (NR) protein sequences from NCBI. Taxonomic information was integrated using DIAMOND’s “makedb” option with the following input files: a protein accession to taxonomy ID mapping file prot.accession2taxid (https://ftp.ncbi.nih.gov/pub/taxonomy/accession2taxid/), a taxonomy nodes file nodes.dmp (https://ftp.ncbi.nih.gov/pub/taxonomy/new_taxdump/), and a taxonomy names file names.dmp (https://ftp.ncbi.nih.gov/pub/taxonomy/new_taxdump/). The processed reads were aligned to the prepared database using DIAMOND’s blastx mode with “very-sensitive” option, ensuring comprehensive taxonomic classification as per the most stringent options of the package. This step translated nucleotide sequences into protein sequences, aligning them against the database. The alignment output format was specified to include detailed taxonomic information for each read. Only the best hit for each read was retained using the option “max-target-seqs 1”. The results were parsed with in-house python script to aggregate reads based on the NCBI taxonomy structure for different phyla of Protostomes and Deuterostomes. Because this reduced version of the CZ ID pipeline was used to overcome the read subset limitations and the exclusive identification of eukaryotes, the contig generation step of the pipeline was not implemented, as it is not feasible to reconstruct species-specific contigs for eukaryotes due to the large genome sizes, the pooled nature and the unequal representation of species in eDNA NGS data. The metazoan metagenomics pipeline has been deposited in GitHub (https://github.com/nousiaso). Assessing our eDNA samples for human reads was conducted with University of Florida Institutional Review Board (IRB-01) ethical approval under project number IRB202201336. Area-proportional Venn diagrams of genus lists were generated using BioVenn^48^.

### qPCR

The Applied Biosystem pre-validated Taqman Gene Expression qPCR assay directed against the human *ZNF285* gene (assay ID Hs00603276_s1) was used as species-specific human assays, based on having no cross reactivity with over 29 other species from mammals to plants (https://www.thermofisher.com/order/genome-database/ and Whitmore et al. 2023^15,19^, Mouse, Rat, Arabidopsis, *C. elegans*, Fruit fly, Bovine, Dog, Chinese hamster, Goat, White-tufted-ear marmoset, Guinea pig, Zebrafish, Horse, Chicken, Soybean, Cynomolgus monkey, Sheep, Rabbit, Green sea turtle, Loggerhead sea turtle, Rice, Rhesus monkey, Baker’s yeast, Fission yeast, Pig, Bread wheat, Wine grape, Western clawed frog and Maize), and having both primers and probe within a single exon (i.e. detects DNA). A pan-eukaryotic 18s rRNA gene (Applied Biosystem, 4319413E) pre-validated Taqman Gene Expression assays, which also has both primers and probe within a single exon (i.e. detects DNA), was used to quantify the total level of pan-eukaryotic DNA in each of the samples. We have previously applied this pan-eukaryotic assay as a positive eDNA extraction and PCR inhibitor control for a range of eDNA sample types^11,13,14,19^.

The qPCR reaction mixtures were performed on 384-well plates in a total volume of 10 μl per well: 5 μl TaqMan Gene Expression Master Mix (Fisher Scientific Cat No: 4369016); 3.5 μl Nuclease free water (Fisher Scientific); 0.5 μl of the respective assay (primer/probe mix, manufacturer supplied concentration); 1 μl DNA template (or, for no-template controls, an additional 1 μl of nuclease-free water per well). Each biological sample was run in 3 technical replicates for eDNA samples and standard curve points. Note that for the previously published qPCR results (2022 samples), 3 technical replicates (18s assay), or 6 technical replicates (ZNF285 assay) were conducted^19^. Negative field controls had the same number of technical replicates run as their corresponding eDNA samples. No template controls were run in triplicate on every qPCR plate. qPCR reactions were performed on an Applied Biosystems QuantStudio 7 Flex, at The Conway Institute’s Genomics Core, University College Dublin, Ireland with the following cycling parameter: 50 °C for 2 min and 95° C for 10 min for one cycle, followed by 45 cycles of 95 °C for 15 s and 60 °C for 1 min. For absolute quantity qPCRs, a standard curve was generated using six DNA one-in-ten serial dilutions, ranging from 111.3 ng/μl to 0.001113 ng/μl. 1 μl of template was used per standard curve reaction. The standard curve template used for both pan-eukaryotic and human specific qPCRs was DNA extracted from cultured human IMR32 neuroblastoma cells in May 2013 for the Duffy et al. (2018) study^46^. For each standard curve concentration and each eDNA sample technical triplicate wells/reactions.

qPCR results were plotted with BoxPlotR^49^ (http://shiny.chemgrid.org/boxplotr/) with every datapoint displayed. Tukey whiskers (extend to data points that are less than 1.5 x IQR away from 1st/3rd quartile) were utilized for every boxplot. One box is graphed per single sample, consisting of all qPCR technical replicate wells for that sample. Biological replicates are denoted by letters A-D at the end of the sample name. Biological replicates are not pooled on any boxplots, with each sample being denoted by its own box.

## Acknowledgements

Sincerest thanks to Paula Murphy and Rebecca Rolfe for facilitating eDNA extraction of Irish samples in their lab in the Zoology Department, Trinity College Dublin, to Aleksandar Krstic, Walter Kolch, Amaya Garcia Munoz and the Conway Core Facilities staff for facilitating qPCR of the samples at Systems Biology Ireland and the Conway Institute of Biomolecular and Biomedical Research at University College Dublin and to BMKGene for conducting the sequencing. Thanks also to Mark Martindale, Nancy Condron, Jennifer Hines and Catherine Eastman.

---

**Supplemental Figure 1.**
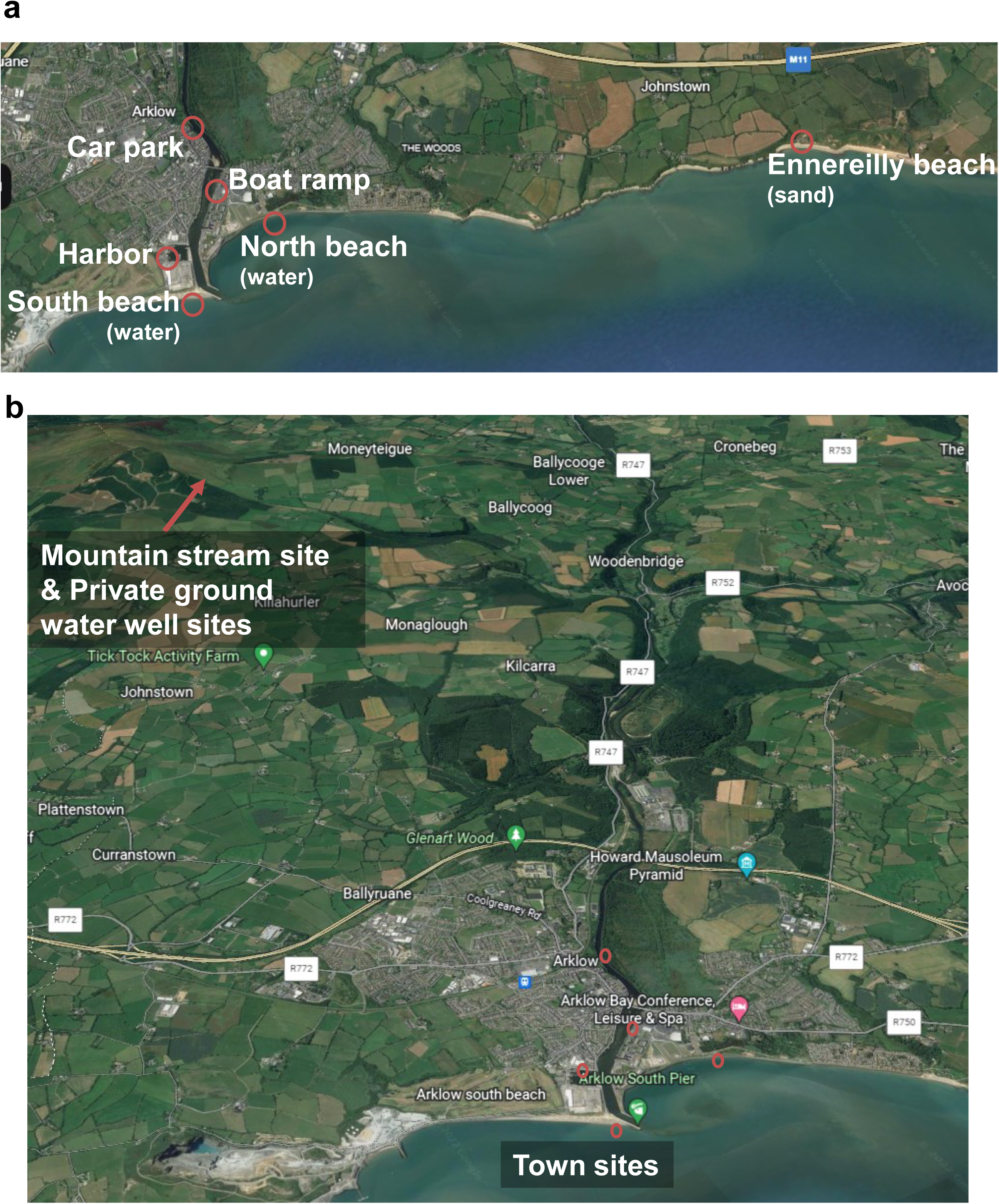
**a)** Satellite imagery view (Google Earth, Map data ©2021 Google and TerraMetrics) showing the locations of the coastal sampling sites, for water (Arklow) and the beach sand (Ennereilly) samples. **b)** Satellite imagery view (Google Earth, Map data ©2021 Google and TerraMetrics) showing the locations of all water sampling sites (Avoca River and Goldmines tributary), adapted from Whitmore et al. 2023^19^.

**Supplemental Figure 2.**
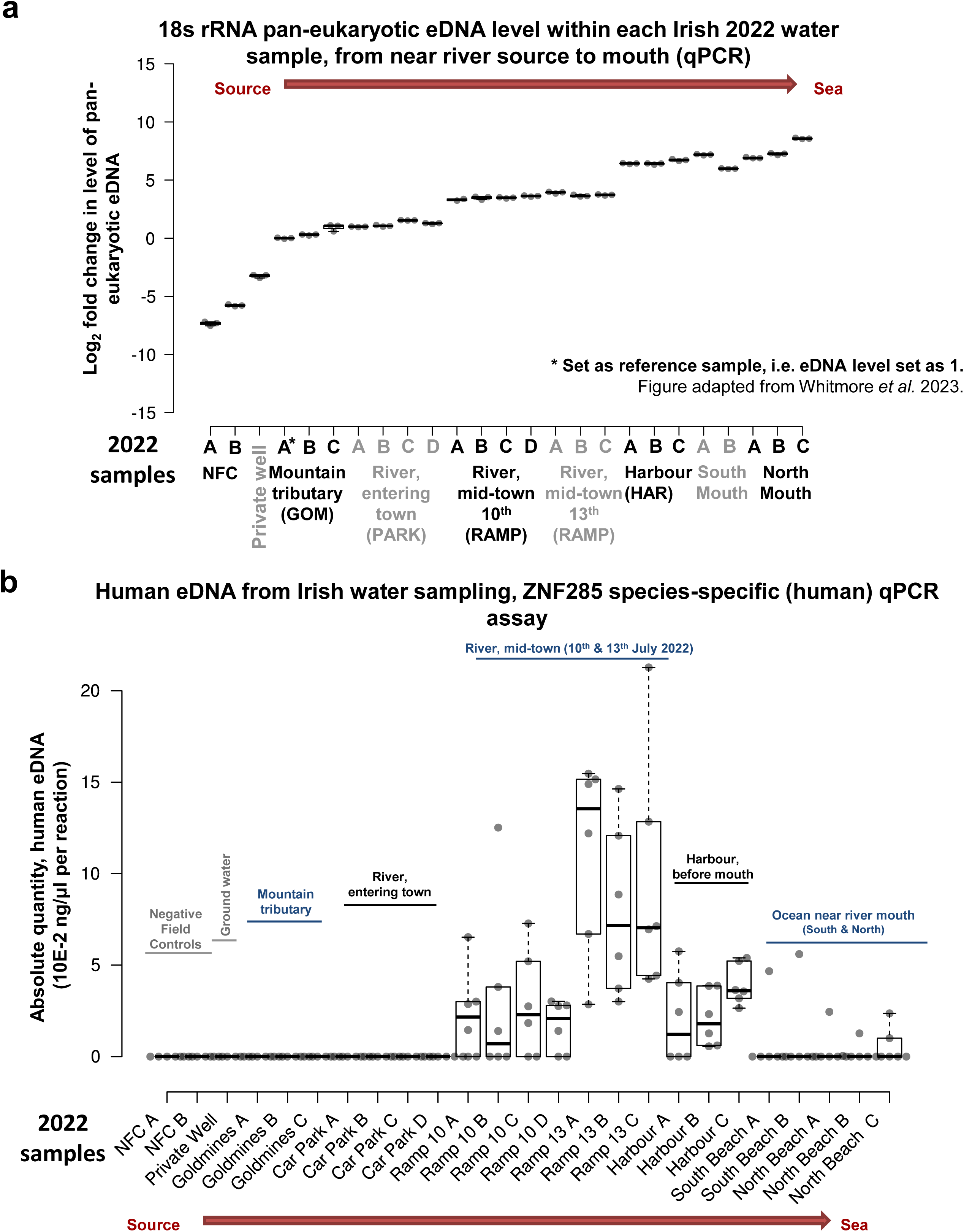
**a)** Pan-eukaryotic eDNA levels within each 2022 water eDNA sample. Relative quantity of 18 s rRNA pan-eukaryotic eDNA levels within each eDNA sample, from near river source to mouth, as assessed by qPCR. Each qPCR reaction is a 10 μl reaction containing 1 μl of extracted eDNA template. For filtered water volumes and elution volumes see Whitmore et al. (2023) and Supplemental Table 1. Figure panel adapted from Whitmore et al. (2023)^19^. **b)** qPCR-based species-specific quantification of human eDNA from Avoca River water sampling, intentional capture. Absolute quantity (10E-2 ng/μl per reaction) of human eDNA per sample. Each qPCR reaction is a 10 μl reaction containing 1 μl of extracted eDNA template. Quantified with *ZNF285* human-specific assay. For filtered water volumes and elution volumes see Whitmore et al. (2023) and Supplemental Table 1. For matching results from these samples quantified with *LILRB2* human-specific assay see Whitmore et al. (2023)^19^. Figure panel adapted from Whitmore et al. (2023)^19^.

**Supplemental Figure 3.**
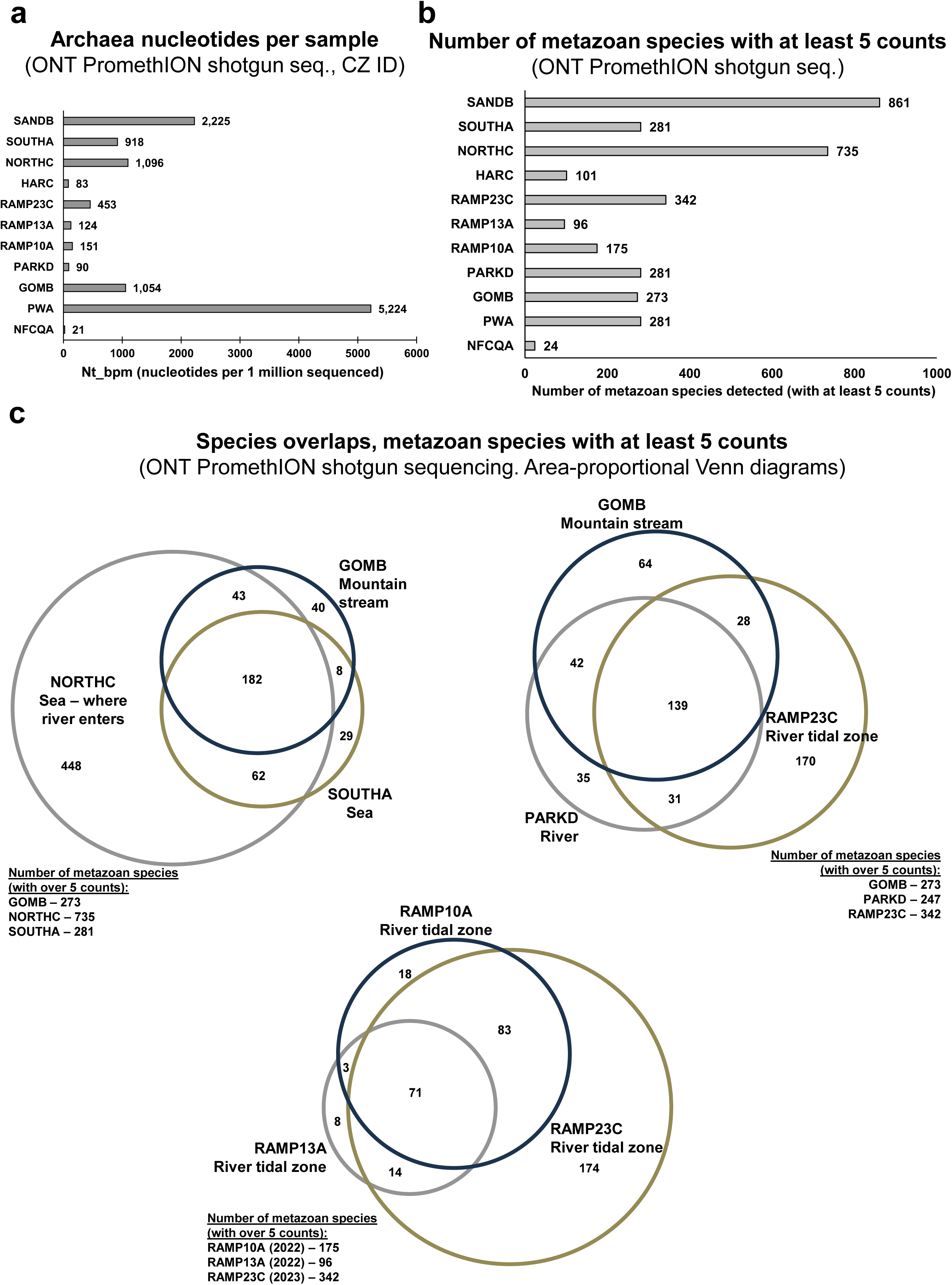
**a)** Quantification of all Archaea hits throughout the sample set, arranged from freshwater to sea water (and finally beach sand), as detected by CZ ID metagenomic analysis of the long-read shotgun sequencing. **b)** Number of metazoan species, with at least 5 read counts, detected by long-read shotgun sequencing in each sample. **c)** Area-proportional Venn diagrams of metazoan species overlaps between eDNA samples, showing the number and overlap of metazoan species with at least 5 counts per sample, as detected by ONT shotgun sequencing. Generated using BioVenn^48^. See Figure 2b for additional sample comparisons.

## References

1. Pfenning-Butterworth, A. et al. Interconnecting global threats: climate change, biodiversity loss, and infectious diseases. The Lancet Planetary Health 8, e270–e283 (2024).

2. Sandbrook, C. et al. Social considerations are crucial to success in implementing the 30×30 global conservation target. Nature Ecology & Evolution 7, 784–785 (2023).

3. Stammnitz, M.R., Hartman Scholz, A. & Duffy, D.J. Environmental DNA without borders. EMBO reports 25, 4095–4099 (2024).

4. Convention_on_Biological_Diversity. Convention on Biological Diversity. COP15: final text of Kunming-Montreal Global Biodiversity Framework. Vol. https://www.cbd.int/article/cop15-final-text-kunming-montreal-gbf-221222 (2022).

5. Aucone, E. et al. Drone-assisted collection of environmental DNA from tree branches for biodiversity monitoring. Science Robotics 8, eadd5762 (2023).

6. Abrego, N. et al. Airborne DNA reveals predictable spatial and seasonal dynamics of fungi. Nature 631, 835–842 (2024).

7. Lynggaard, C. et al. Vertebrate environmental DNA from leaf swabs. Current Biology 33, R853–R854 (2023).

8. Aalismail, N.A., Díaz-Rúa, R., Geraldi, N., Cusack, M. & Duarte, C.M. Diversity and sources of airborne eukaryotic communities (AEC) in the global dust belt over the Red Sea. Earth Systems and Environment 5, 459–471 (2021).

9. Urban, L. et al. Freshwater monitoring by nanopore sequencing. eLife 10, e61504 (2021).

10. Farrell, J.A., Whitmore, L. & Duffy, D.J. The promise and pitfalls of environmental DNA and RNA approaches for the monitoring of human and animal pathogens from aquatic sources. BioScience 71, 609–625 (2021).

11. Farrell, J.A. et al. Environmental DNA monitoring of oncogenic viral shedding and genomic profiling of sea turtle fibropapillomatosis reveals unusual viral dynamics. Communications Biology 4, 565 (2021).

12. Bass, D., Christison, K.W., Stentiford, G.D., Cook, L.S.J. & Hartikainen, H. Environmental DNA/RNA for pathogen and parasite detection, surveillance, and ecology. Trends in Parasitology 39, 285–304 (2023).

13. Koda, S.A. et al. A novel eDNA approach for rare species monitoring: Application of long-read shotgun sequencing to Lynx rufus soil pawprints. Biological Conservation 287, 110315 (2023).

14. Farrell, J.A. et al. Detection and population genomics of sea turtle species via non-invasive environmental DNA analysis of nesting beach sand tracks and oceanic water. Molecular Ecology Resources 22, 2471–2493 (2022).

15. McCauley, M., Koda, S.A., Loesgen, S. & Duffy, D.J. Multicellular species environmental DNA (eDNA) research constrained by overfocus on mitochondrial DNA. Science of The Total Environment 912, 169550 (2024).

16. Kjær, K.H. et al. A 2-million-year-old ecosystem in Greenland uncovered by environmental DNA. Nature 612, 283–291 (2022).

17. Djurhuus, A. et al. Environmental DNA reveals seasonal shifts and potential interactions in a marine community. Nature Communications 11, 254 (2020).

18. Crits-Christoph, A. et al. Genetic tracing of market wildlife and viruses at the epicenter of the COVID-19 pandemic. Cell 187, 5468–5482.e11 (2024).

19. Whitmore, L. et al. Inadvertent human genomic bycatch and intentional capture raise beneficial applications and ethical concerns with environmental DNA. Nature Ecology & Evolution 7, 873– 888 (2023).

20. Marx, V. Method of the year: long-read sequencing. Nature Methods 20, 6–11 (2023).

21. Broman, E., Bonaglia, S., Norkko, A., Creer, S. & Nascimento, F.J.A. High throughput shotgun sequencing of eRNA reveals taxonomic and derived functional shifts across a benthic productivity gradient. Molecular Ecology 30, 3023–3039 (2021).

22. Bista, I. et al. Performance of amplicon and shotgun sequencing for accurate biomass estimation in invertebrate community samples. Molecular Ecology Resources 18, 1020–1034 (2018).

23. Downie, A.T., Bennett, W.W., Wilkinson, S., de Bruyn, M. & DiBattista, J.D. From land to sea: Environmental DNA is correlated with long-term water quality indicators in an urbanized estuary. Marine Pollution Bulletin 207, 116887 (2024).

24. Perry, W.B. et al. An integrated spatio-temporal view of riverine biodiversity using environmental DNA metabarcoding. Nature Communications 15, 4372 (2024).

25. Agustinho, D.P. et al. Unveiling microbial diversity: harnessing long-read sequencing technology. Nature Methods 21, 954–966 (2024).

26. Liem, M., Regensburg-Tuïnk, T., Henkel, C., Jansen, H. & Spaink, H. Microbial diversity characterization of seawater in a pilot study using Oxford Nanopore Technologies long-read sequencing. BMC Research Notes 14, 42 (2021).

27. Benítez-Páez, A. & Sanz, Y. Multi-locus and long amplicon sequencing approach to study microbial diversity at species level using the MinION™ portable nanopore sequencer. GigaScience 6(2017).

28. Lewin, H.A. et al. Earth BioGenome Project: Sequencing life for the future of life. Proceedings of the National Academy of Sciences 115, 4325–4333 (2018).

29. Nakano, H. What is Xenoturbella? Zoological Letters 1, 22 (2015).

30. Bourlat, S.J. et al. Deuterostome phylogeny reveals monophyletic chordates and the new phylum Xenoturbellida. Nature 444, 85–88 (2006).

31. Simmonds, S.E., et al. CZ ID: a cloud-based, no-code platform enabling advanced long read metagenomic analysis. *bioRxiv*, 2024.02.29.579666 (2024).

32. Fisher, M.C. & Garner, T.W.J. Chytrid fungi and global amphibian declines. Nature Reviews Microbiology 18, 332–343 (2020).

33. Scheele, B.C. et al. Amphibian fungal panzootic causes catastrophic and ongoing loss of biodiversity. Science 363, 1459–1463 (2019).

34. Venter, J.C. et al. Environmental genome shotgun sequencing of the Sargasso Sea. Science 304, 66–74 (2004).

35. Duffy, D.J. Problems, challenges and promises: perspectives on precision medicine. Briefings in Bioinformatics 17, 494–504 (2016).

36. Twyford, A. et al. A DNA barcoding framework for taxonomic verification in the Darwin Tree of Life Project [version 1; peer review: 2 approved]. Wellcome Open Research 9(2024).

37. Guo, M., Yuan, C., Tao, L., Cai, Y. & Zhang, W. Life barcoded by DNA barcodes. Conservation Genetics Resources 14, 351–365 (2022).

38. Antil, S. et al. DNA barcoding, an effective tool for species identification: a review. Molecular Biology Reports 50, 761–775 (2023).

39. Kalantar, K.L. et al. IDseq—An open source cloud-based pipeline and analysis service for metagenomic pathogen detection and monitoring. GigaScience 9(2020).

40. Ahmed, W. et al. First confirmed detection of SARS-CoV-2 in untreated wastewater in Australia: A proof of concept for the wastewater surveillance of COVID-19 in the community. Science of The Total Environment 728, 138764 (2020).

41. Hart, O.E. & Halden, R.U. Computational analysis of SARS-CoV-2/COVID-19 surveillance by wastewater-based epidemiology locally and globally: Feasibility, economy, opportunities and challenges. Science of The Total Environment 730, 138875 (2020).

42. Zan, R. et al. Environmental DNA clarifies impacts of combined sewer overflows on the bacteriology of an urban river and resulting risks to public health. Science of The Total Environment 889, 164282 (2023).

43. Ahmed, W., Hughes, B. & Harwood, V.J. Current Status of Marker Genes of Bacteroides and Related Taxa for Identifying Sewage Pollution in Environmental Waters. Water 8, 231 (2016).

44. Nagler, M., Podmirseg, S.M., Ascher-Jenull, J., Sint, D. & Traugott, M. Why eDNA fractions need consideration in biomonitoring. Molecular Ecology Resources 22, 2458–2470 (2022).

45. Seymour, M. et al. Acidity promotes degradation of multi-species environmental DNA in lotic mesocosms. Communications Biology 1, 4 (2018).

46. Duffy, D.J. et al. Identification of a MYCN and Wnt-related VANGL2-ITLN1 fusion gene in neuroblastoma. Gene Reports 12, 187–200 (2018).

47. Buchfink, B., Xie, C. & Huson, D.H. Fast and sensitive protein alignment using DIAMOND. Nature Methods 12, 59–60 (2015).

48. Hulsen, T., de Vlieg, J. & Alkema, W. BioVenn - a web application for the comparison and visualization of biological lists using area-proportional Venn diagrams. BMC Genomics 9, 488 (2008).

49. Spitzer, M., Wildenhain, J., Rappsilber, J. & Tyers, M. BoxPlotR: a web tool for generation of box plots. Nature Methods 11, 121–122 (2014).

